# DiffRhyShiny: A Benchmarking-Informed Platform for Differential Rhythmicity Analysis with Dependence-Aware Evidence Integration

**DOI:** 10.64898/2026.10.07.757323

**Authors:** Chuan Liu, Lu Liu, Na Li, Yang Jiang, Chunyan Li, Ying Liu, Yongan Fu, Kai Guo

## Abstract

**Motivation:** Differential rhythmicity analysis identifies condition-specific changes in biological oscillations, but diverse algorithmic assumptions complicate method selection. Furthermore, integrating complementary evidence across different tools while accounting for statistical dependence remains an unresolved challenge.

**Results:** We benchmarked seven algorithms across diverse rhythmic scenarios, revealing that optimal selection depends heavily on specific biological variations and experimental designs. To overcome single-algorithm limitations, we introduced Kost’s Method, a dependence-aware *p*-value integration approach. Building on this framework, we developed DiffRhyShiny, an interactive web platform that automates robust differential rhythmicity analysis, consensus evaluation, and visualization. Furthermore, we validated the platform’s performance and statistical rigor using both extensive simulated scenarios and real-world transcriptomic datasets.

**Availability and implementation:** DiffRhyShiny is freely available at https://omicsai.shinyapps.io/diffrhyshiny, with the latest source code at https://github.com/liuchuan111/diffrhyshiny. The code and data to reproduce all benchmarking results are archived on Figshare at https://doi.org/10.6084/m9.figshare.33070100.

## Introduction

High-throughput technologies have expanded chronobiology from the analysis of individual clock-controlled genes to genome-wide investigations of rhythmic molecular processes (Wu G *et al*. 2014). Current circadian transcriptomic studies use bulk RNA sequencing, proteomics and metabolomics, and more recently single-cell approaches (Mukherji *et al*. 2024, Xu *et al*. 2024, Zhu *et al*. 2024, Qin *et al*. 2025), to characterize thousands of rhythmic features across biological conditions. Circadian dysfunction is implicated in a wide range of diseases, making comparisons of rhythmic patterns between biological conditions, including healthy and disease states, different tissues, and environmental contexts, an important analytical goal (Zha *et al*. 2025). This analytical task is referred to as differential rhythmicity analysis, which aims to identify changes in rhythmic properties between groups rather than simply detect rhythmic signals within a single condition. However, selecting appropriate computational methods remains challenging because available algorithms rely on different statistical assumptions and evaluate distinct aspects of rhythmic variation (Parsons *et al*. 2020).

Several computational methods have been developed for differential rhythmicity analysis, including DODR, LimoRhyde, CircaCompare, DiffCircaPipeline, compareRhythms, dryR, DiffCircadian, and RepeatedCircadian (Thaben and Westermark 2016, Singer and Hughey 2019, Ding *et al*. 2021, 2022, Weger *et al*. 2021, Pelikan *et al*. 2022, Xue *et al*. 2023). Among these approaches, seven representative methods with compatible computational frameworks and available implementations were selected for benchmarking in this study. These algorithms use different statistical frameworks and define differential rhythmicity from distinct perspectives. For example, DODR estimates overall rhythmic changes through robust regression (Thaben and Westermark 2016), whereas LimoRhyde applies likelihood ratio testing to evaluate rhythmic differences (Singer and Hughey 2019). CircaCompare, diffCircadian, and DiffCircaPipeline compare fitted rhythmic models and parameters between conditions (Parsons *et al*. 2020, Ding *et al*. 2021, Xue *et al*. 2023). In contrast, compareRhythms and dryR rely on Bayesian model selection to classify rhythmic patterns (Weger *et al*. 2021, Pelikan *et al*. 2022). Due to differences in model assumptions, input requirements, and statistical criteria, these methods may identify different sets of differential rhythmic genes from the same dataset. However, the extent to which their performance varies under different biological scenarios and experimental designs remains insufficiently characterized.

Previous studies have compared differential rhythmicity algorithms and demonstrated substantial disagreement among their results. Miao et al. reported that different algorithms generated distinct differential rhythmic gene sets from the same transcriptomic dataset, largely because each method applies different statistical definitions of rhythmic alteration (Miao *et al*. 2024). Several software platforms have also been developed to improve accessibility of circadian analysis tools (Wu B and Ja 2025, Han *et al*. 2026). However, important methodological questions remain unresolved. First, no comprehensive benchmark has measured how accurately each algorithm classifies the specific type of rhythmic change present, whether both conditions are rhythmic with a shared phase, or one is arrhythmic, or phases differ. Second, the influence of unequal noise levels between experimental conditions has not been systematically investigated, despite its potential impact on statistical inference. Third, the effects of sampling frequency, observation duration, biological replication, and missing observations on algorithm performance remain unclear.

Combining results from multiple algorithms represents another unresolved challenge. In single-condition analysis, MetaCycle integrates multiple *p*-values using Fisher’s method; however, this approach assumes independence among tests, an assumption that may not hold when multiple statistical tests are applied to the same gene expression profile (Wu G *et al*. 2016). Kost’s Method provides a dependence-aware alternative by incorporating correlations among *p*-values during statistical integration (Kost and McDermott 2002). Although this approach has been applied in other bioinformatics contexts, its application to differential rhythmicity analysis has not been evaluated in this study area. Here, we systematically benchmark seven differential rhythmicity methods across multiple simulation scenarios and transcriptomic datasets. We further introduce a Kost’s Method-based consensus framework and implement it in DiffRhyShiny, an interactive platform that integrates algorithm execution, statistical comparison, and visualization within a unified workflow.

## Materials and Methods

### RNA-seq Data Acquisition and Preprocessing

We benchmarked the methods using the mouse fibroblast dataset GSE253826 from the Gene Expression Omnibus. This dataset contains RPKM-normalized time-series profiles from wild-type and Bmal1 knockout fibroblasts and has previously been used for circadian and differential rhythmicity analyses (Miao *et al*. 2024, Unruh *et al*. 2024). The design comprised 24 time points across the two conditions.

Circadian rhythm genes MetaCycle with minper=24, maxper=24, and cycMethod=c(“ARS”, “JTK”, “LS”). Genes with a Benjamini–Hochberg adjusted <0.25 in either condition were retained, yielding 866 genes for subsequent analyses. The benchmark included DODR, LimoRhyde/LimoRhyde2, CircaCompare, diffCircadian, DiffCircaPipeline, compareRhythms, and dryR.

### Algorithmic Configurations and Parameters

All methods were evaluated using RPKM-normalized expression values (Mortazavi *et al*. 2008). Although LimoRhyde2, compareRhythms, and dryR can accept raw counts, a common normalized input was used to compare algorithmic performance independently of preprocessing differences.

Methods were grouped into hypothesis-testing approaches, including DODR, LimoRhyde, CircaCompare, diffCircadian, and DiffCircaPipeline, and model-selection approaches, including dryR and compareRhythms. The period was fixed at 24 h. Raw p-values were adjusted using the Benjamini–Hochberg procedure, whereas methods with intrinsic adjusted metrics used their default significance criteria. Software versions and settings are provided in Table 1.

**Table 1.**
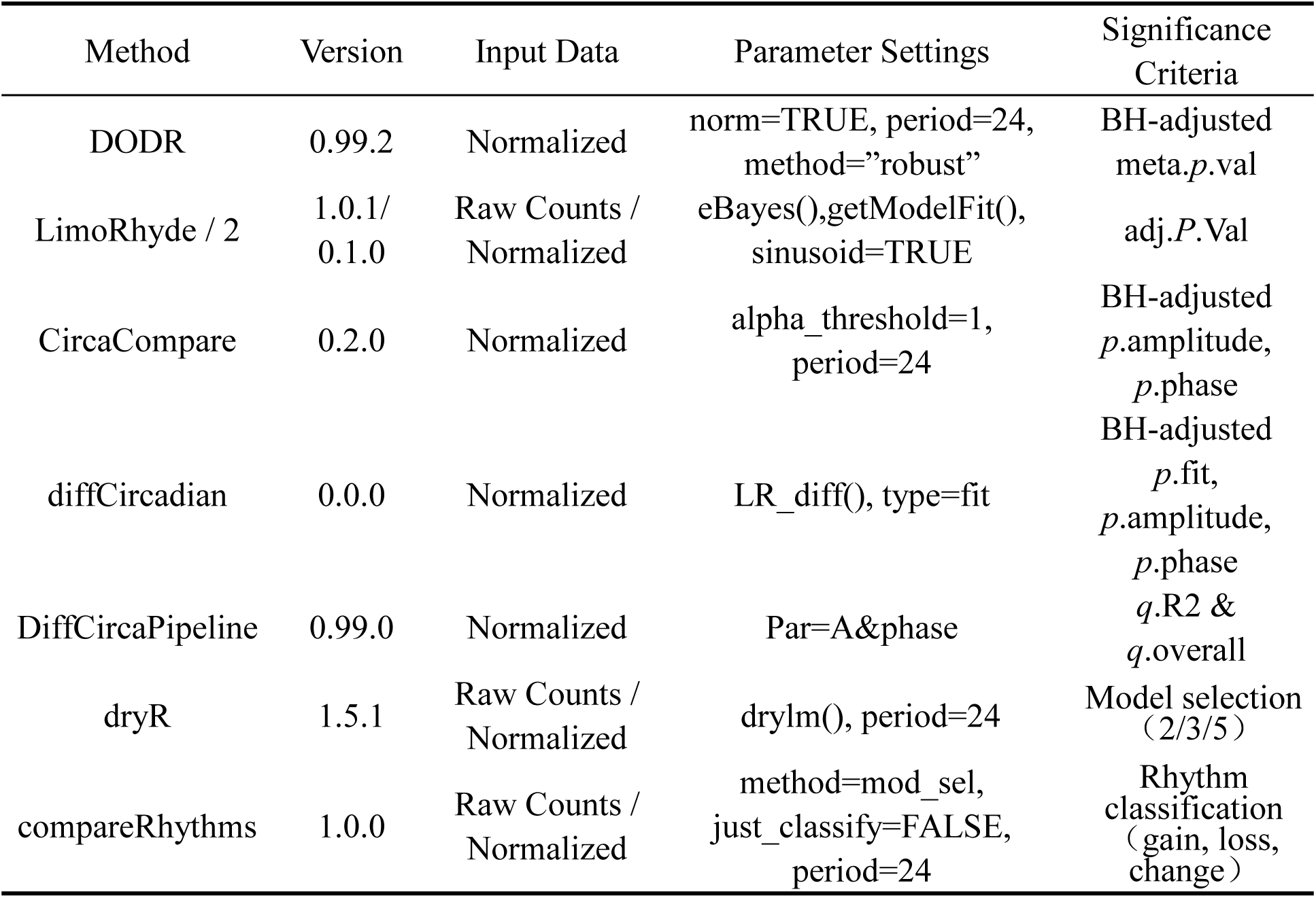
Parameter settings and selection criteria for different algorithms.

### Generation of Simulated Data for Classification

For classification benchmarking, we simulated two groups sampled at 0, 4, 8, 12, 16, and 20 h, with five replicates per time point and 30 samples per group. Gene expression profiles were generated using a cosine model:

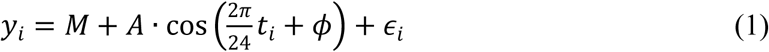

Where A represents amplitude, ɸis phase, M is the mesor (mid-line estimating statistic of rhythm), and €_i_N∼(0, *σ*^2^) is Gaussian noise.

Five scenarios were grouped into three ground-truth classes: Arrhy, with A_1_ = A _2_ = 0; Rhy I, with A_1_ > 0 and A_2_ = 0 across varying amplitudes; and Both rhythmic. The latter included Both-same, with identical parameters; Both-different phase, with _1_ ≠ _2_; and Both-different SNR, with identical rhythmic parameters but SNR_1_ ≠ SNR_2_.

Six algorithms capable of single-condition rhythmic classification or parameter-based comparison were evaluated: DiffCircaPipeline, compareRhythms, dryR, CircaCompare, diffCircadian, and LimoRhyde. DODR was excluded from this analysis because it does not provide individual rhythmic parameter estimates or direct classification outputs required for this evaluation.

Direct classifications were retained when available. Otherwise, genes were labeled Arrhy when neither group was rhythmic, Rhy I or Rhy II when only one group was rhythmic, and Both when both groups were rhythmic, using *p*<0.05. Accuracy was the proportion correctly classified within each category and was averaged across 10 replicates.

### Generation of Simulated Data with Differential Parameters

To estimate parameter-specific power, we simulated two rhythmic groups differing only in mesor, amplitude, or phase. Each group contained 30 observations sampled randomly within 0 to 24 h, with expression generated using Eq. (1). The rhythmic parameters of Group I were fixed as:

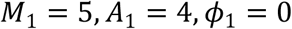

Differential rhythmicity was introduced by modifying only one parameter in Group II while maintaining the remaining parameters unchanged.

Three types of parameter alterations were simulated:

i. Mesor variation: Amplitude and phase were fixed:

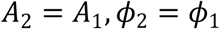

while mesor values were varied:

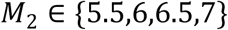

(ii) Phase variation: Mesor and amplitude were fixed:

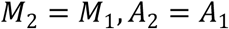

while phase shifts were introduced:

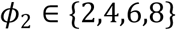

(iii) Amplitude variation: Mesor and phase were fixed:

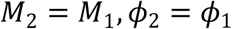

while amplitude values were varied:

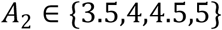

The noise level was maintained at a constant value across all simulation conditions to isolate the effect of rhythmic parameter changes. For each parameter combination, 10,000 genes were simulated and divided into ten independent replicates. Statistical power was calculated as the proportion of truly differential rhythmic genes correctly identified by each algorithm.

### Simulation of Noise Heterogeneity

To assess noise heterogeneity, we defined = /, where is amplitude and is the Gaussian noise standard deviation. We evaluated Type I error under equal noise levels and power when groups had identical rhythmic parameters but unequal SNRs.

For the null condition, both groups were simulated with identical rhythmic parameters:

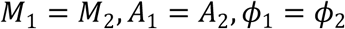

Therefore, no differential rhythmicity was present, and significant detections were considered false positives. For each parameter combination, 10,000 genes were simulated. Sampling times were randomly generated from the interval [0,24] h. Noise levels were adjusted to produce SNR values of 0.4, 0.6, 0.8, 1.0, 1.2, 1.5, 2.0, 2.5, 3.0, 3.5, and 4.0, covering a range of rhythmic signal strengths reported in mammalian circadian transcriptomic studies (Xue *et al*. 2023).

For the alternative condition, rhythmic parameters were maintained between groups:

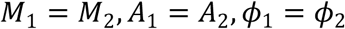

while noise levels were independently varied between groups to generate different SNR values:

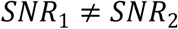

This simulation design evaluated whether differential rhythmicity algorithms could distinguish noise heterogeneity from changes in underlying rhythmic parameters. Algorithm performance was assessed using Type I error rates under the null condition and statistical power under the alternative condition.

### Generation of Multi-scenario Simulated Datasets

For broader detection and sampling benchmarks, we generated cosine-based datasets because CircaCompare, diffCircadian, and DiffCircaPipeline use sinusoidal parameterization. Eight rhythmic categories were grouped as two evaluation classes:

Non-differential rhythmic genes (Non-DRGs) included Arrhy and Same, with 3,000 genes per category. DRGs included Rhy I, Rhy II, diffA, diffPhase, diffM, and diffSNR, with 1,000 genes per category. The dataset therefore contained equal numbers of DRGs and Non-DRGs.

### Evaluation of Sampling Strategies

To investigate the influence of experimental design on algorithm performance, we simulated different sampling strategies by varying sampling frequency, observation duration, biological replication, and missing observations. Sampling density was evaluated by comparing three 24-h sampling schemes: 1. 4h/1d: Six time points collected at 4-h intervals. 2. 3h/1d: Eight time points collected at 3-h intervals. 3. 2h/1d: Twelve time points collected at 2-h intervals.

We further evaluated the effect of extending the observation window to 48 h while maintaining comparable numbers of observations. To investigate the trade-off between sampling frequency and biological replication, experimental designs with different allocations of sampling points and replicates were compared under similar total sample sizes.

To simulate incomplete longitudinal measurements, one, five, or eight time points were randomly removed from the standard 4h/1d sampling scheme. Algorithm performance was then evaluated across these sampling conditions using classification accuracy.

### Architecture and Statistical Framework of DiffRhyShiny

DiffRhyShiny was developed as an interactive web platform using the R Shiny framework to integrate multiple differential rhythmicity algorithms into a unified analysis workflow. The platform adopts a modular user interface and server architecture, with the frontend implemented using shinydashboard to support workflow navigation and parameter configuration. The backend integrates seven differential rhythmicity algorithms: DODR, LimoRhyde/LimoRhyde2, CircaCompare, diffCircadian, DiffCircaPipeline, compareRhythms, and dryR. Because these algorithms differ in input requirements and output formats, DiffRhyShiny incorporates preprocessing modules implemented using the R packages data.table and tidyr for data formatting, sample annotation, and output harmonization. Interactive visualization components based on Plotly enable users to examine rhythmic expression profiles, fitted curves, statistical results, and algorithm comparisons within the same analytical environment.

To integrate statistical evidence generated by multiple differential rhythmicity algorithms, DiffRhyShiny implements Kost’s Method for dependent *p*-value combination (Kost and McDermott 2002). This approach was selected because different algorithms analyze the same gene expression profiles and therefore may produce correlated statistical evidence. For each gene, let _1_, _2_, …, represent the *p*-values generated from statistical tests derived from different algorithms or rhythmic parameters. For example, CircaCompare provides separate statistical tests for amplitude and phase differences, whereas diffCircadian evaluates multiple components including model fit, amplitude, and phase. Kost’s Method combines these correlated *p*-values using a scaled chi-square distribution, where the effective degrees of freedom are adjusted according to the estimated covariance structure among input statistics (Kost and McDermott 2002). Unlike Fisher’s method, which assumes independence among combined tests, Kost’s Method accounts for dependency among statistical evidence sources during *p*-value integration.

Kost’s Method was implemented using the EmpiricalBrownsMethod Bioconductor package (version 1.34.0), which provides an empirical approximation for dependent *p*-value combination. For algorithms generating raw *p*-values, the original *p*-values were used as inputs for integration. For DODR, which primarily reports the BH-adjusted meta.*p*.val statistic, the available raw *p*-value was used when provided; otherwise, the reported adjusted value was used consistently across analyses. The resulting consensus statistic was interpreted as a combined measure of evidence from multiple algorithms rather than a replacement for individual algorithm outputs. The consensus framework was evaluated through three complementary approaches: case-level analysis using the mouse fibroblast transcriptomic dataset (GSE253826), simulation-based benchmarking using datasets with predefined ground-truth labels, and independent biological evaluation using the liver feeding-rhythm dataset (GSE135898). DiffRhyShiny was deployed through the shinyapps.io (https://omicsai.shinyapps.io/diffrhyshiny/) platform to provide browser-based access, and the source code was made publicly available for local deployment and further development.

## Results

### Quantitative Heterogeneity and Consensus Patterns of DRGs

To characterize differences among differential rhythmicity algorithms, we quantified the number of differential rhythmic genes (DRGs) identified by each method using the *Bmal1* knockout transcriptomic dataset (GSE253826) across multiple statistical thresholds (Fig. 1A). The number of detected DRGs varied substantially among algorithms, reflecting differences in statistical models and significance criteria. Among the evaluated methods, diffCircadian identified the largest number of DRGs across the tested q-value thresholds, whereas DiffCircaPipeline showed a more conservative detection profile, particularly under the stringent threshold of BH-adjusted *q*-value < 0.05. DODR and LimoRhyde showed comparable detection patterns for global rhythmicity analysis, while the Bayesian model-selection approaches, dryR and compareRhythms, produced similar numbers of identified DRGs.

**Fig. 1.**
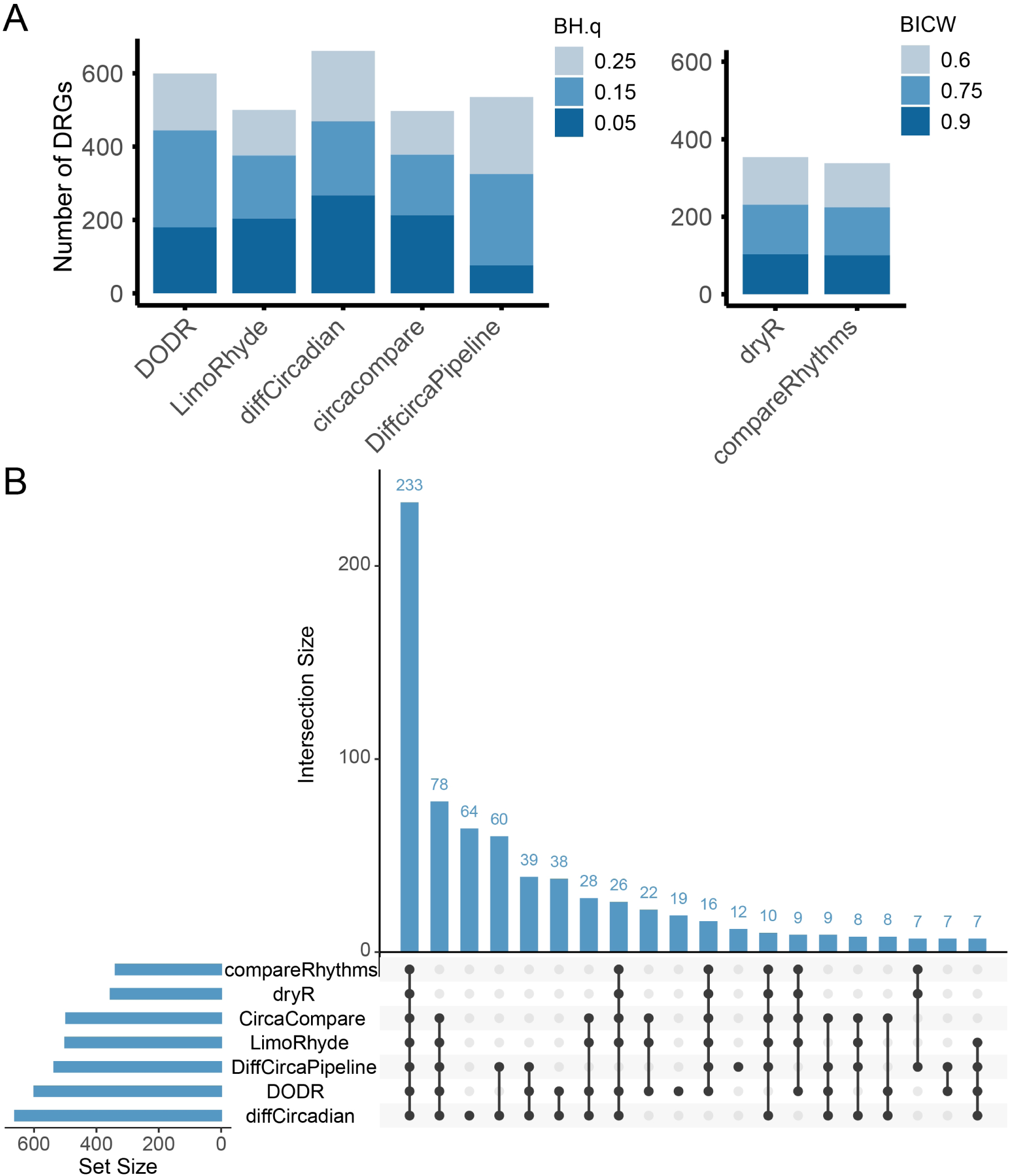
DRG identification and overlap across algorithms. **(A)** Sensitivity across statistical thresholds (BH.*q* values for significance-test methods, left; BIC weights for Bayesian methods, right). **(B)** UpSet plot showing the intersection of identified DRG signatures. Alt text: A bar chart and an UpSet plot displaying the number of differentially rhythmic genes and their overlap identified by seven different algorithms across various statistical thresholds.

To further evaluate the agreement among algorithms, we performed an intersection analysis of DRGs identified by the seven methods (Fig. 1B). A total of 233 genes were detected by all algorithms, representing the shared DRG set across different analytical frameworks. However, each algorithm also identified method-specific gene sets. For example, diffCircadian detected 64 additional DRGs that were not identified by the other evaluated methods. The overlap patterns between algorithms demonstrated substantial differences in candidate gene selection despite analysis of the same transcriptomic dataset. These findings indicate that algorithm choice influences the number and composition of detected DRGs.

### Consistency of Rhythmic Parameter Estimation Among Algorithms

To evaluate agreement among algorithms in estimating rhythmic parameters, we compared amplitude and phase changes among the DRGs identified by all seven methods. We focused on the 233 shared DRGs detected by all algorithms to evaluate whether different computational frameworks produced comparable estimates for the same genes. Six algorithms that provide numerical rhythmic parameter estimates were included: LimoRhyde2, CircaCompare, diffCircadian, DiffCircaPipeline, compareRhythms, and dryR.

Because these algorithms report rhythmic changes using different output formats, including direct parameter differences or condition-specific estimates, all results were converted into a unified control-minus-knockout representation before correlation analysis. Pearson correlation coefficients were then calculated to evaluate agreement among methods for amplitude and phase changes. The results showed strong positive correlations among all evaluated algorithms for both amplitude and phase estimates, with Pearson correlation coefficients greater than 0.928 across the 233 shared DRGs (Fig. 2). CircaCompare, DiffCircaPipeline, and diffCircadian showed correlations approaching 1.0 for parameter estimates, while compareRhythms and dryR also demonstrated strong agreement with each other. LimoRhyde2 showed slightly lower correlation coefficients, ranging from 0.928 to 0.935, but remained positively correlated with the other methods. These results indicate that although differential rhythmicity algorithms may differ in DRG identification, they produce highly consistent rhythmic parameter estimates for genes detected by all methods.

**Fig. 2.**
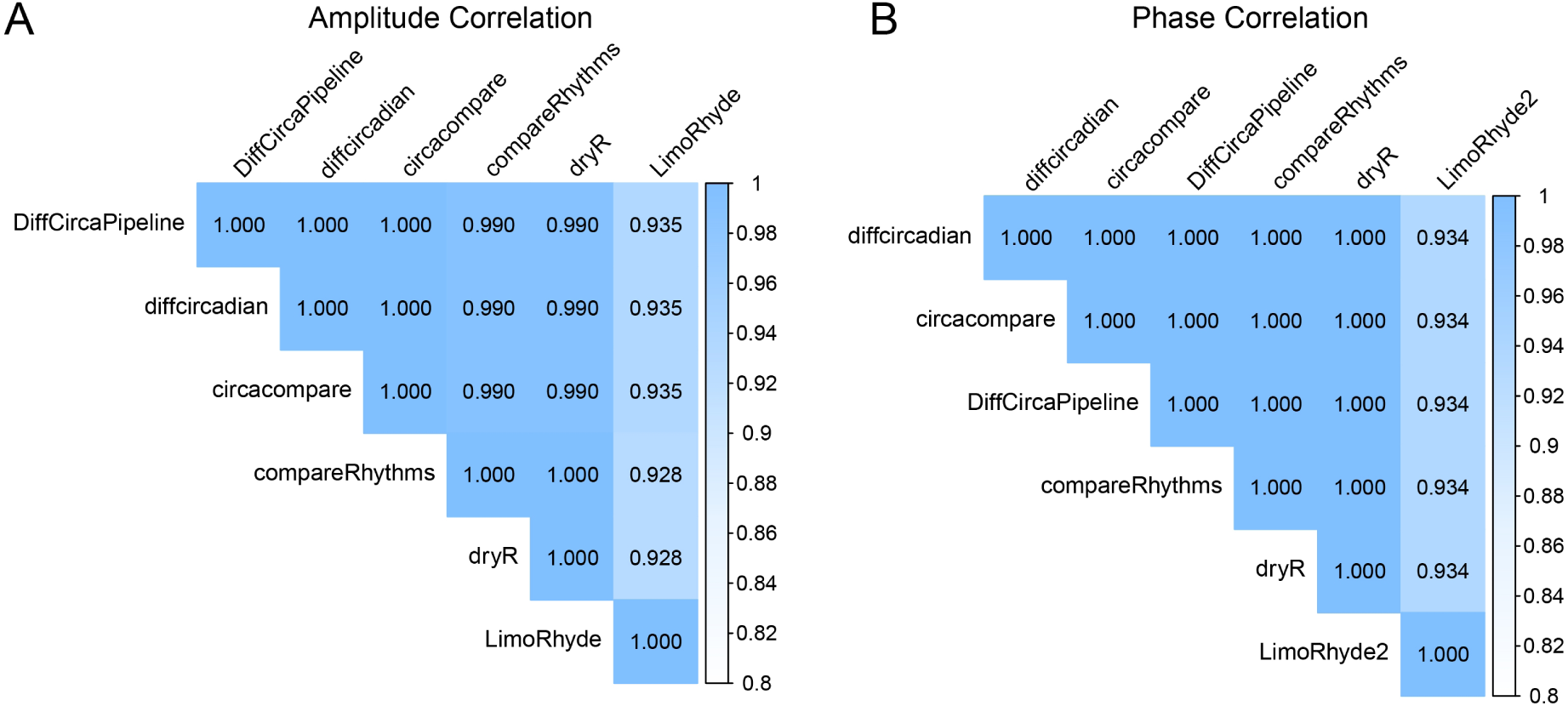
Consistency of rhythmic parameter estimation. Pairwise Pearson correlation heatmaps of **(A)** amplitude shifts and **(B)** phase shifts calculated from the 233 consensus DRGs. Alt text: Two heatmaps illustrating the pairwise Pearson correlation coefficients among different algorithms for estimating amplitude and phase shifts of consensus genes.

### Benchmarking Algorithm Classification Performance Across Rhythmic Scenarios

To evaluate the ability of differential rhythmicity algorithms to classify distinct rhythmic patterns, we generated five simulated scenarios: Arrhythmic, Rhythmic in Group I only (Rhy I), Both-same, Both-different phase, and Both-different SNR. These scenarios represented different combinations of rhythmic presence, rhythmic parameter changes, and noise variation between groups. Because DODR does not provide single-group rhythmic classification or individual rhythmic parameter estimates required for this analysis, it was excluded from this evaluation. The remaining six algorithms, including DiffCircaPipeline, compareRhythms, dryR, CircaCompare, diffCircadian, and LimoRhyde, were compared using classification accuracy.

Using the predefined classification criteria based on single-group rhythmic significance (< 0.05), the evaluated algorithms showed different performance patterns across simulation scenarios (Fig. 3). For the Arrhythmic category, DiffCircaPipeline achieved the highest classification accuracy (0.953), whereas other algorithms showed lower accuracy under the same condition (Fig. 3A). For the Rhy I scenario, CircaCompare showed increasing classification accuracy as the amplitude difference between groups increased (Fig. 3B).

**Fig. 3.**
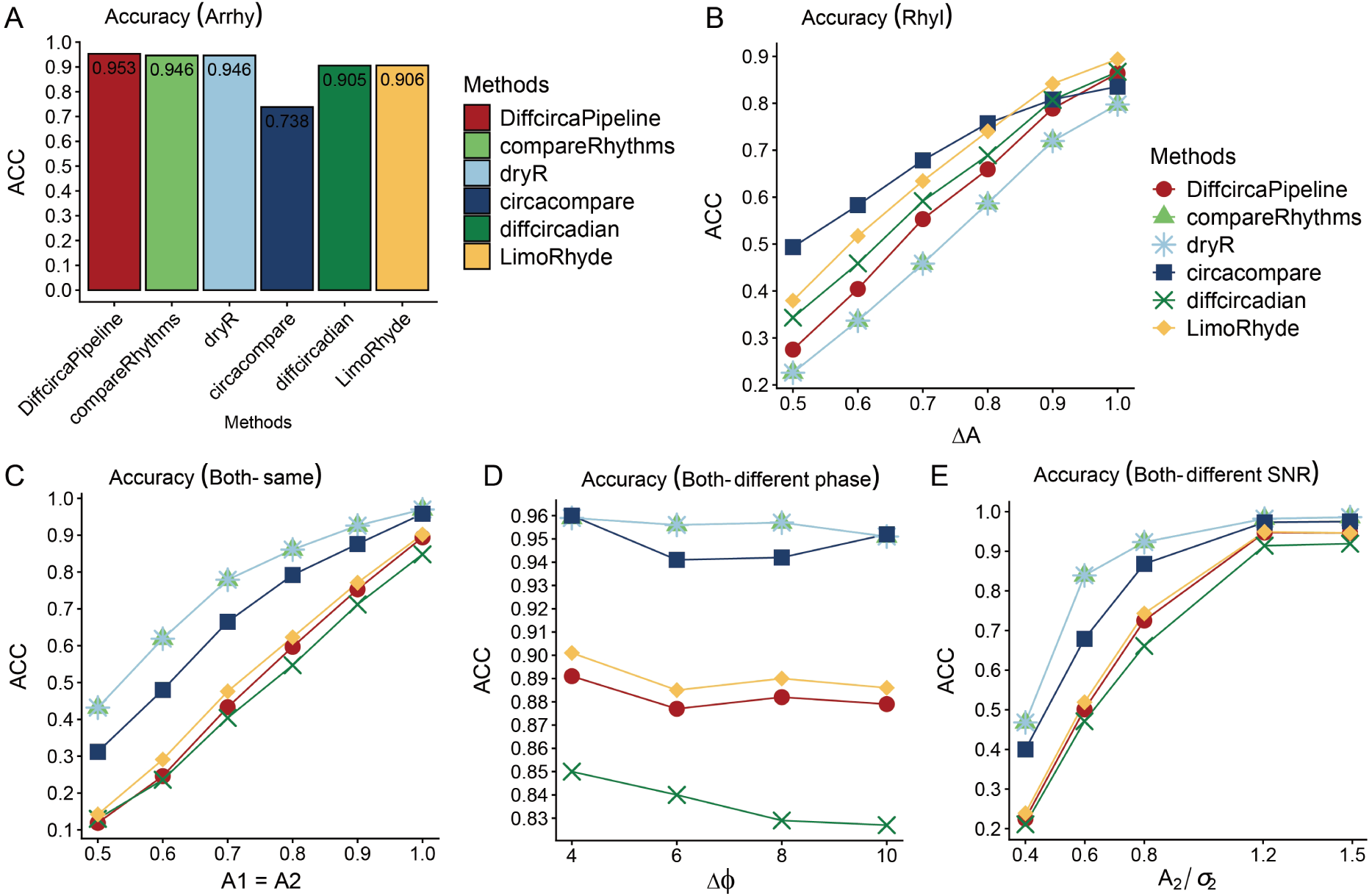
Algorithmic classification accuracy across rhythmic scenarios. Accuracy (ACC) of six algorithms evaluated across simulated categories: **(A)** Arrhythmic (Arrhy) patterns for noise filtration; **(B)** Rhythmic in Group I (RhyI) patterns across an amplitude gradient; and **(C–E)** dual-rhythmic (Both-class) scenarios featuring **(C)** identical parameters, **(D)** phase variations, or **(E)** Signal-to-Noise Ratio (SNR) differences. Data points represent the mean of ten replicates. Alt text: Line and bar graphs evaluating the classification accuracy of six algorithms across five simulated rhythmic scenarios, including varying amplitudes, phases, and signal-to-noise ratios.

For scenarios in which both groups were rhythmic, model-selection-based approaches showed higher classification accuracy than several hypothesis-testing approaches (Fig. 3C–E). When both groups shared identical rhythmic parameters at a moderate amplitude (_1_ = _2_ = 0.8), dryR and compareRhythms correctly classified 86.1% of genes as non-differential rhythmic genes. In comparison, diffCircadian and DiffCircaPipeline achieved accuracies of 54.7% and 59.7%, respectively (Fig. 3C).

The performance similarity between certain algorithms was also observed across simulation categories. compareRhythms and dryR showed highly similar classification patterns, while diffCircadian and LimoRhyde, as well as DiffCircaPipeline and LimoRhyde in specific scenarios, showed comparable results (Fig. 3). Overall, no single algorithm achieved the highest accuracy across all simulated conditions. Instead, algorithm performance depended on the specific rhythmic scenario evaluated.

### Benchmarking Type I Error Control and Power for Rhythmic Parameter Detection

To evaluate Type I error control and statistical power, we generated simulated datasets with predefined rhythmic parameter structures. Under the null condition, where no differences in rhythmic parameters existed between groups, we assessed whether algorithms maintained error rates close to the nominal significance level of 0.05 (Fig. 4A). DiffCircaPipeline, CircaCompare, diffCircadian, and LimoRhyde maintained Type I error rates close to the expected level under both dual-parameter and triple-parameter settings. In contrast, DODR showed increased false-positive rates under the evaluated simulation conditions. DODR was retained in subsequent analyses as a representative method for comparison with other differential rhythmicity approaches.

**Fig. 4.**
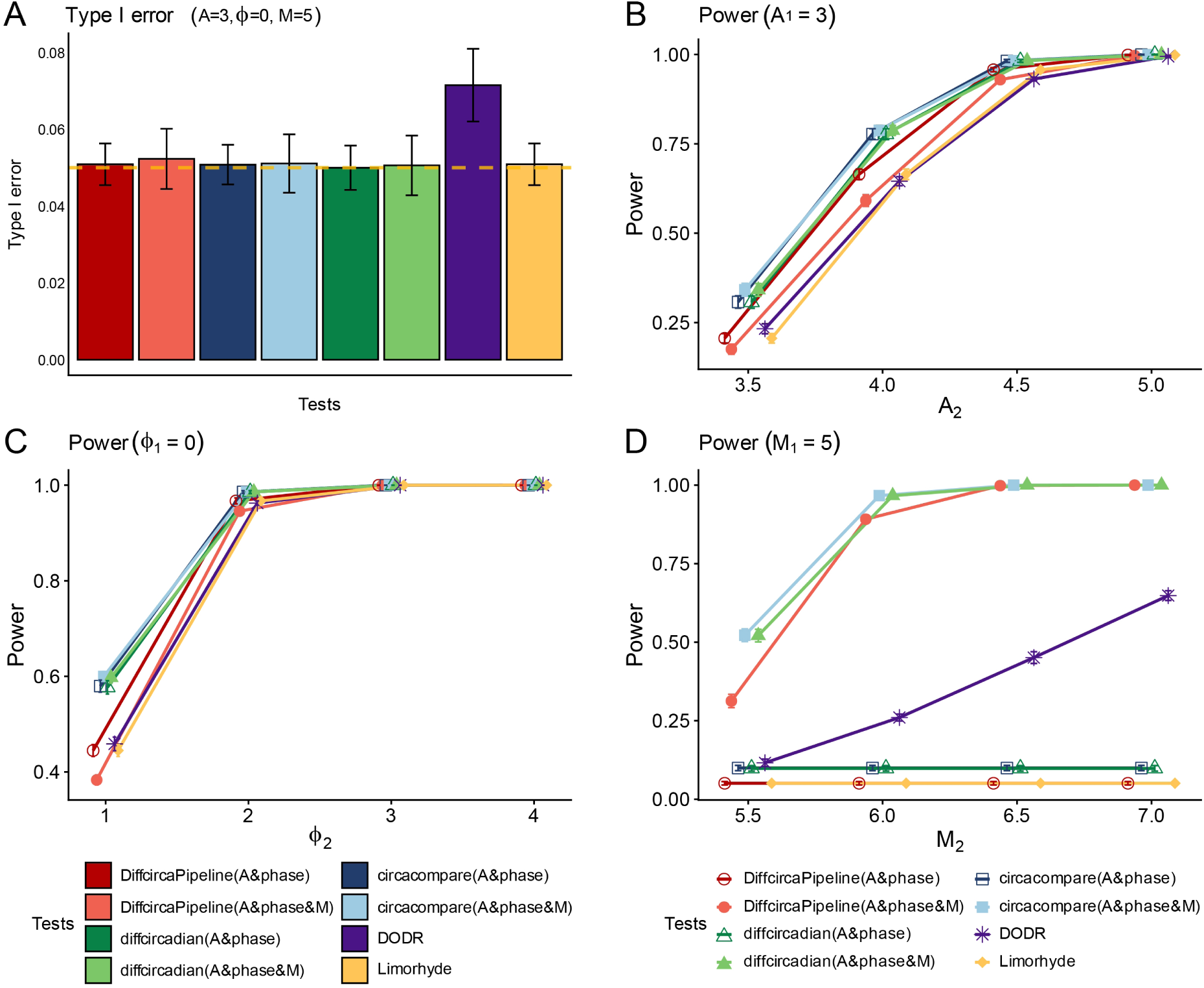
Statistical power and Type I error control in detecting rhythmic parameter shifts. **(A)** Type I error rates under null-hypothesis conditions (= 3, = 0, = 5; dashed line: nominal 0.05 level). **(B–D)** Statistical power in detecting shifts in **(B)** amplitude, **(C)** phase, and **(D)** Mesor. Data points represent the mean of ten replicates; error bars indicate SD. Alt text: Bar charts and line graphs showing the Type I error rates and statistical power of different algorithms in detecting simulated shifts in amplitude, phase, and mesor.

Model-selection-based methods, including compareRhythms and dryR, were not included in this parameter-specific power analysis because they classify rhythmic patterns using Bayesian Information Criterion weights rather than directly testing individual rhythmic parameter differences.

Under alternative simulation conditions, all evaluated algorithms showed increasing statistical power as the magnitude of rhythmic parameter differences increased. For amplitude and phase variation scenarios, dual-parameter models generally achieved higher power than corresponding triple-parameter models (Fig. 4B,C). CircaCompare achieved the highest statistical power among the evaluated methods in these two scenarios, followed by diffCircadian and DiffCircaPipeline.

The detection performance differed in mesor variation scenarios (Fig. 4D). Algorithms or model configurations that did not include mesor as a tested parameter showed limited power for detecting mesor differences. In contrast, triple-parameter configurations of CircaCompare, diffCircadian, and DiffCircaPipeline showed increased power for identifying mesor alterations. These results demonstrate that algorithm performance depends on whether the tested statistical model includes the rhythmic parameter associated with the simulated biological change.

### Benchmarking Algorithm Performance Under Noise Heterogeneity

To evaluate algorithm performance under unequal noise levels between experimental conditions, we generated two simulation scenarios based on signal-to-noise ratio (SNR) variation: a null condition for assessing Type I error control and an alternative condition for evaluating detection power under noise heterogeneity (Fig. 5).

**Fig. 5.**
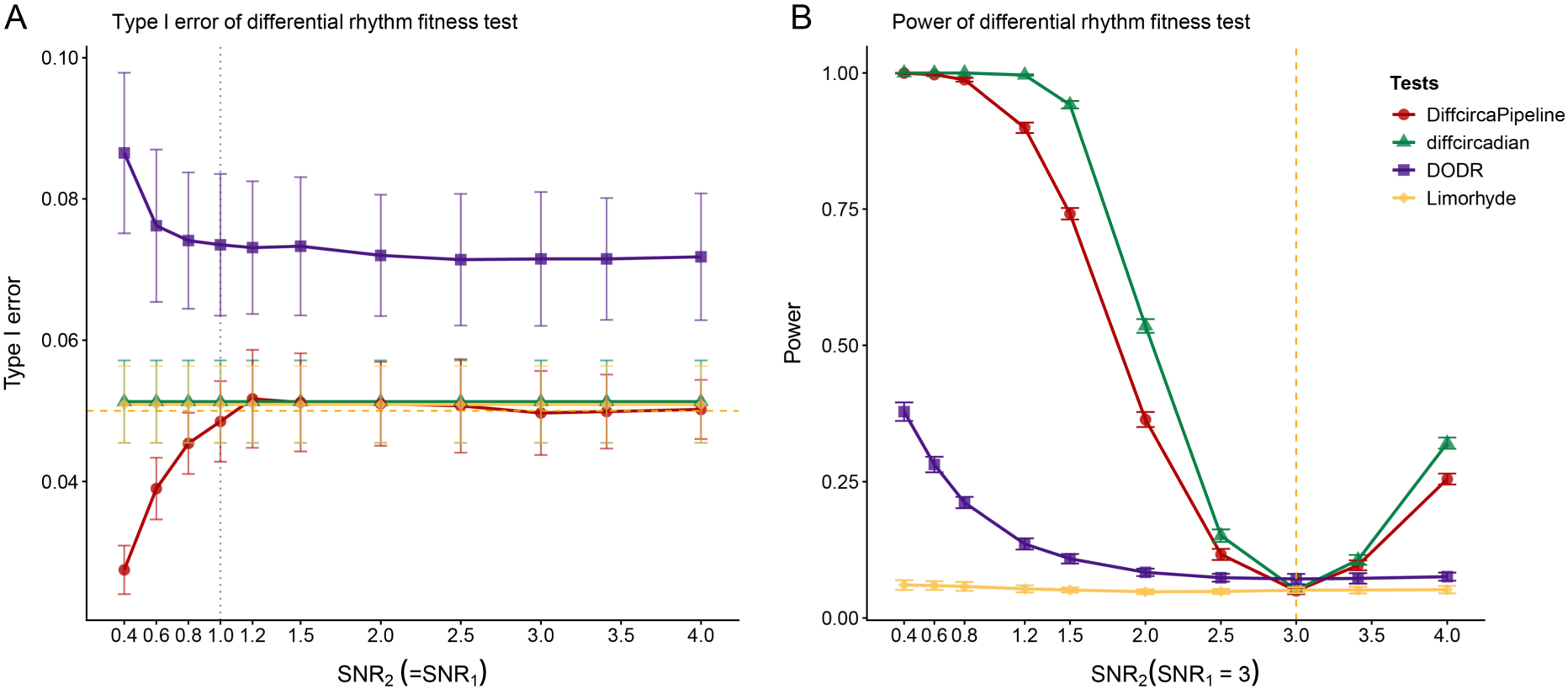
Robustness and sensitivity of algorithms to noise heterogeneity. **(A)** Type I error rates across a gradient of matching SNRs (dashed line: 0.05 threshold). **(B)** Statistical power in detecting noise-induced fit disparities (_1_ fixed at 3 and _2_ varied; vertical line: identical SNR). Data points represent the mean ± SD of ten replicates. Alt text: Line graphs demonstrating the Type I error rates and statistical detection power of algorithms under conditions of varying and heterogeneous signal-to-noise ratios between experimental groups.

Under the null condition, where both groups shared identical rhythmic parameters, LimoRhyde and diffCircadian maintained Type I error rates close to the nominal level across the evaluated SNR range. DiffCircaPipeline showed a more conservative error profile, with lower false-positive rates under high-noise conditions. In contrast, DODR exhibited elevated Type I error rates compared with the expected significance threshold (Fig. 5A).

Under the alternative condition, where rhythmic parameters remained identical but noise levels differed between groups, DiffCircaPipeline and diffCircadian showed increasing statistical power as the SNR difference between groups increased, resulting in a V-shaped performance pattern across the tested conditions (Fig. 5B). LimoRhyde showed relatively limited variation in power across SNR differences, whereas DODR displayed a decreasing power trend under the evaluated conditions.

Together, these results demonstrate that algorithm performance can vary substantially when experimental groups differ in noise characteristics. Methods that incorporate goodness-of-fit-related information showed different response patterns compared with methods that primarily evaluate rhythmic parameter differences under the tested simulation framework.

### Impact of Sampling Strategies on Detection Performance

To evaluate how experimental design influences differential rhythmicity analysis, we simulated different sampling strategies by varying sampling frequency, observation duration, biological replication, and missing data patterns (Fig. 6). Algorithm performance was compared across these sampling conditions using classification accuracy. When sampling schemes contained the same total number of observations, extending the observation window or increasing sampling frequency produced comparable performance under the evaluated conditions (Fig. 6A). Increasing the total number of observations improved classification accuracy across algorithms. However, the magnitude of improvement differed among methods. DiffCircaPipeline showed relatively stable performance under reduced sampling conditions, whereas DODR and diffCircadian showed greater performance variation when sampling points were reduced (Fig. 6B).

**Fig. 6.**
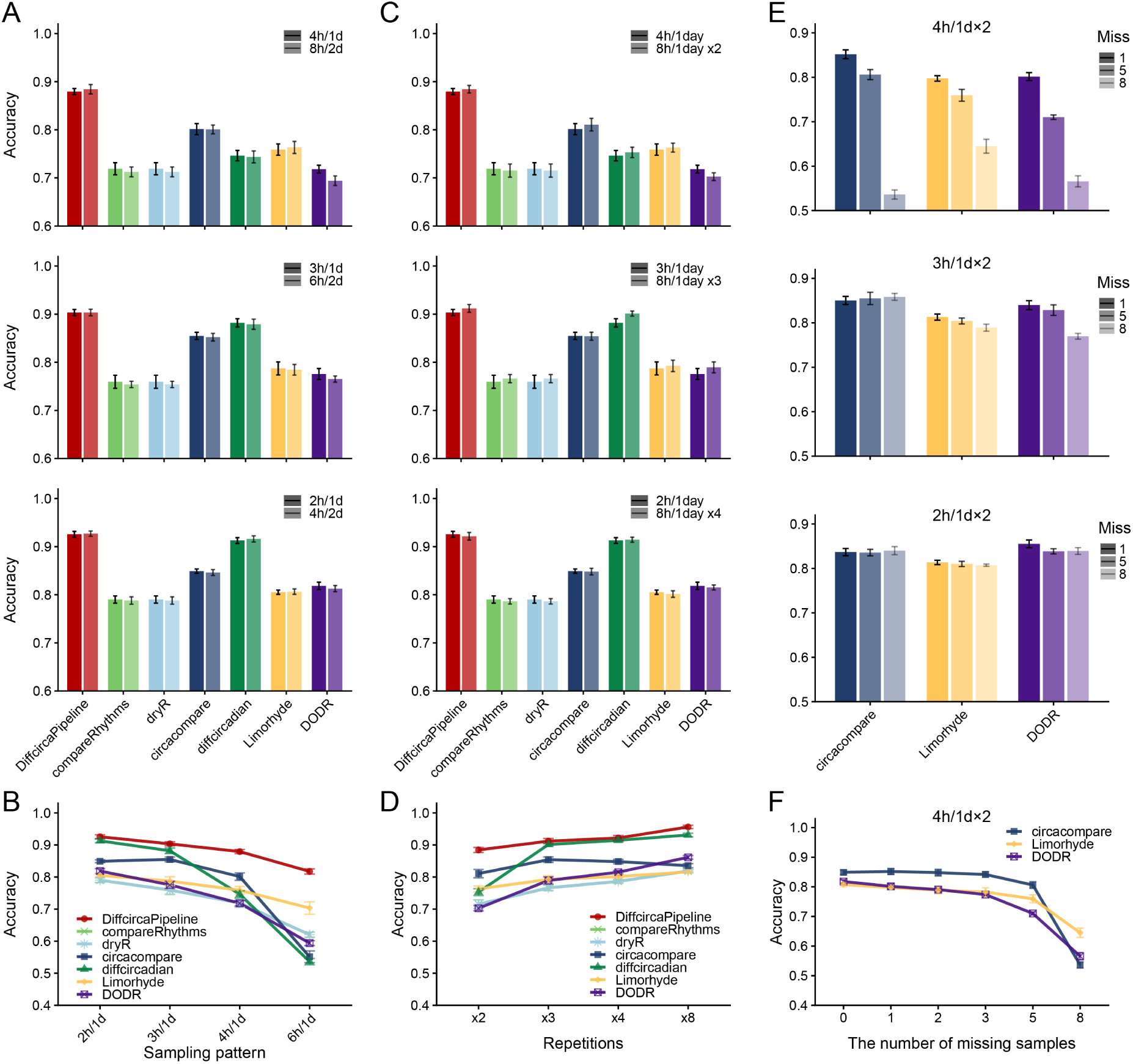
Impact of sampling strategies and data missingness. **(A–B)** Sampling frequency and duration effects comparing **(A)** paired designs with equal observations (e.g., 4h/1d vs. 8h/2d) and **(B)** accuracy decay from 2h/1d to 6h/1d density. **(C–D)** Biological replication vs. density. **(C)** Comparison between single-replicate high-frequency and multi-replicate lower-frequency designs; **(D)** Impact of replication scale (2 to 8). **(E–F)** Tolerance to stochastic data loss showing **(E)** missingness impacts across densities and **(F)** accuracy decay curves (4h/1d×2 scheme). Data represent mean ± SD of ten replicates. Alt text: Bar and line graphs comparing the classification accuracy of algorithms under different sampling frequencies, biological replication scales, and proportions of missing data.

We further evaluated the effect of balancing sampling frequency and biological replication. Several algorithms, including CircaCompare, diffCircadian, and LimoRhyde, achieved higher accuracy in designs with biological replication compared with high-frequency designs containing fewer replicates but similar total numbers of observations (Fig. 6C,D). DODR showed different performance patterns under low and high data-volume conditions, whereas DiffCircaPipeline maintained relatively consistent accuracy across different replication levels. To evaluate tolerance to incomplete time-series data, we randomly removed one, five, or eight time points from the standard 4h/1d sampling scheme (Fig. 6E,F). Classification accuracy decreased as the number of missing observations increased under the standard sampling design.

Higher sampling density partially reduced the performance loss caused by missing observations. Under dense sampling conditions, CircaCompare maintained relatively stable classification accuracy after removal of eight time points.

These results were obtained from controlled cosine-based simulations. Empirical downsampling of real transcriptomic time-series datasets was not performed in this study. Therefore, the influence of sampling design on algorithm performance requires further evaluation using additional experimental datasets.

### DiffRhyShiny: An Integrative Platform for Differential Rhythmicity Analysis

We developed DiffRhyShiny (https://omicsai.shinyapps.io/diffrhyshiny/) as an interactive web platform for differential rhythmicity analysis that integrates seven algorithms, consensus analysis, and visualization modules into a unified workflow (Fig. 7). The platform accepts time-series transcriptomic matrices using a standardized format, where rows represent genes and columns represent samples organized according to group, time point, and biological replicate. DiffRhyShiny provides a three-step workflow consisting of data upload, algorithm execution, and result visualization. The platform includes an algorithm compatibility matrix to assist users in selecting appropriate methods based on analytical objectives. Users can configure analysis parameters, including significance thresholds and classification strategies, and visualize fitted rhythmic curves, statistical summaries, and algorithm comparison results.

**Fig. 7.**
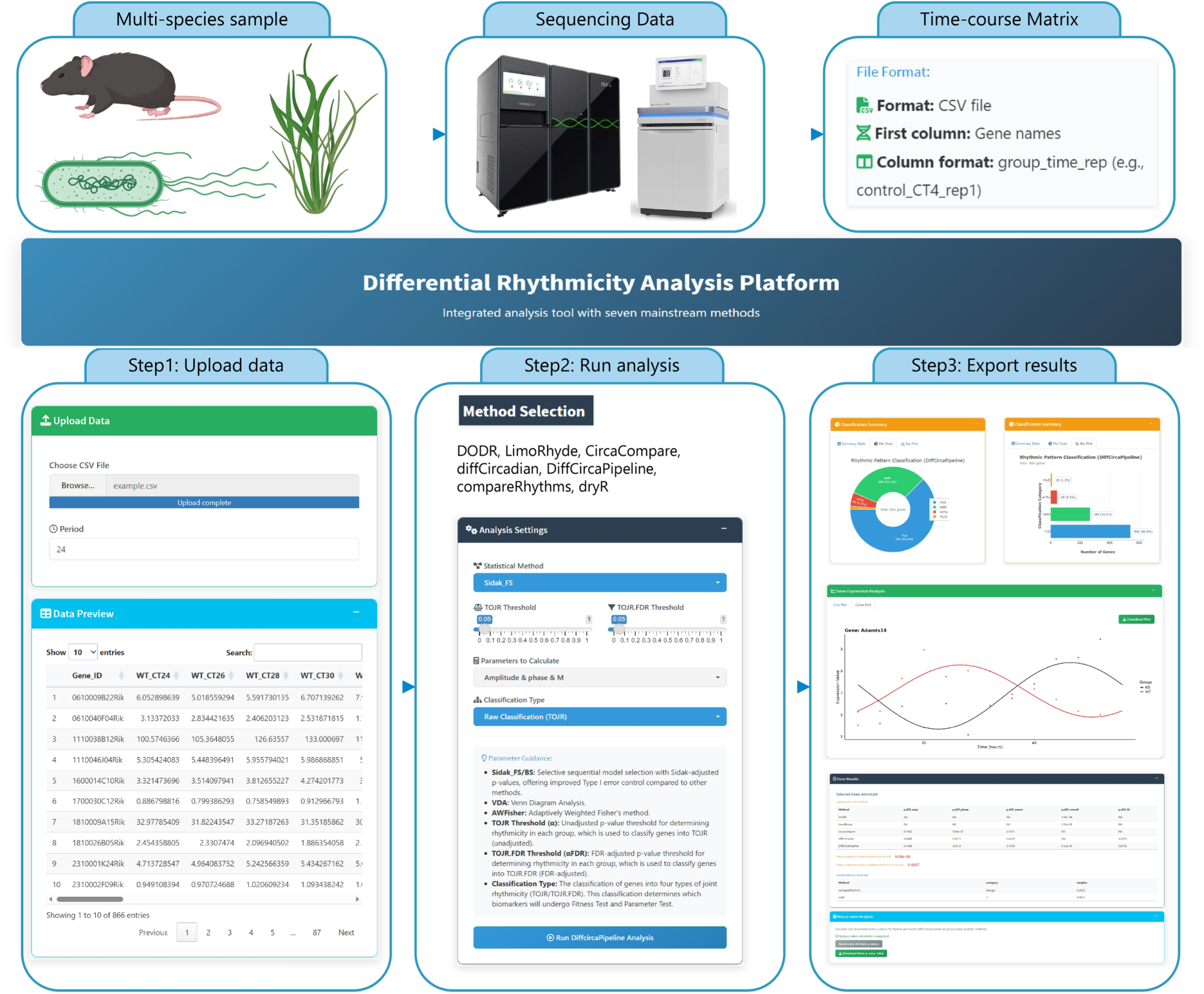
Functional architecture and workflow of the DiffRhyShiny platform. Top panel: Schema of multi-species sample input, sequencing compatibility, and standardized time-course matrix format. Step 1 (Upload data): Graphical interface for CSV uploading, time-period configuration, and matrix preview. Step 2 (Run analysis): Execution dashboard for method selection among seven benchmarked tools and parameter tuning. Step 3 (Export results): Multi-modal exploration dashboards and downstream summary tables for interactive evaluation, exemplified by the Meta *p*-value and Bayesian validation of the phase-shifted *Adamts14* gene. Species cartoons were downloaded from the Generic Diagramming Platform (Jiang *et al*. 2025). Alt text: A schematic flowchart detailing the three-step architecture and user interface of the DiffRhyShiny web platform, from data upload to algorithm execution and interactive results visualization.

To integrate evidence from multiple differential rhythmicity algorithms, DiffRhyShiny implements Kost’s Method for dependent *p*-value combination. Instead of applying a simple intersection of algorithm outputs, the framework combines correlated *p*-values generated from different statistical tests. For a given gene, Kost’s Method estimates covariance among input *p*-values and combines them using a scaled chi-square distribution with adjusted degrees of freedom. This approach accounts for dependency among statistical tests, whereas Fisher’s method assumes independence among combined *p*-values.

We evaluated the consensus framework using three complementary analyses. First, we examined its performance using the *Bmal1* knockout transcriptomic dataset (GSE253826). DiffRhyShiny reproduced the 233 DRGs identified by the manual benchmark analysis (Fig. 1B). The Adamts14 gene provided an example of complementary evidence obtained from multiple algorithms. CircaCompare detected a significant phase difference for Adamts14 (= 1.93 × 10^−7^), whereas amplitude and mesor differences were not significant (> 0.44). Bayesian model selection classified Adamts14 as a rhythmicity-changed gene with a posterior probability of 0.9852. After integration using Kost’s Method, Adamts14 obtained a significant consensus Meta *p*-value of 9.08 × 10^−6^.

Second, we evaluated the statistical properties of the consensus framework using simulated datasets with predefined ground-truth labels. Kost’s Method was compared with Fisher’s method, Brown’s method, and Tippett’s minimum-p approach using *p*-values generated from multiple differential rhythmicity algorithms (Fig. 8). Under null and alternative conditions, the different *p*-value combination methods showed distinct patterns of Type I error control and detection power. Kost’s Method maintained Type I error rates close to the nominal level under the evaluated simulation conditions and showed higher detection performance than more conservative combination strategies under several evaluated alternative scenarios.

**Fig. 8.**
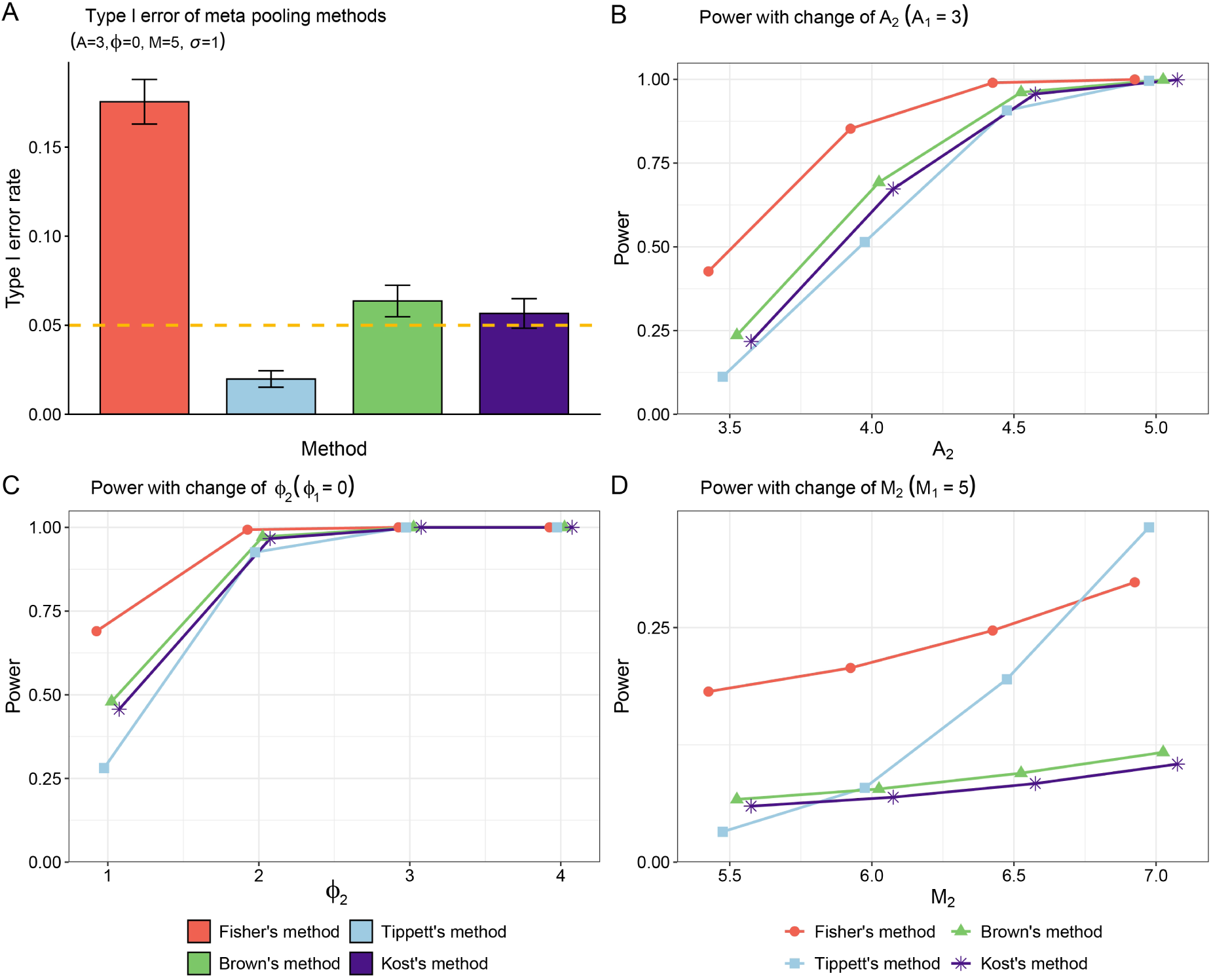
Performance evaluation of dependent-adjusted consensus methods. Benchmarking of four *p*-value combinations (Fisher’s, Tippett’s, Brown’s, and Kost’s) under simulation. **(A)** Empirical Type I error rates under the null hypothesis (A=3, ϕ=0, M=5, =1; dashed line: nominal =0.05). **(B–D)** Empirical power curves across scenarios varying in **(B)** amplitude, **(C)** phase, and **(D)** baseline mesor. Data represent mean ± SD of ten replicates. Alt text: Line graphs comparing empirical Type I error rates and statistical power curves among Fisher’s, Tippett’s, Brown’s, and Kost’s *p*-value combination methods across various simulated differential rhythmicity scenarios.

Third, we evaluated the consensus framework using an independent liver feeding-rhythm dataset (GSE135898). The analysis compared Meta *p*-value results with intersection-based consensus and dryR-based classification. Using dryR-defined food-driven differential rhythmic genes as a reference set, the intersection approach identified 12.62% of these genes, whereas the Meta *p*-value approach identified 24.31% (Fig. 9A,B). These results indicate that dependence-aware *p*-value integration can identify additional genes overlapping with a biologically defined differential rhythmicity signature compared with strict intersection-based approaches.

**Fig. 9.**
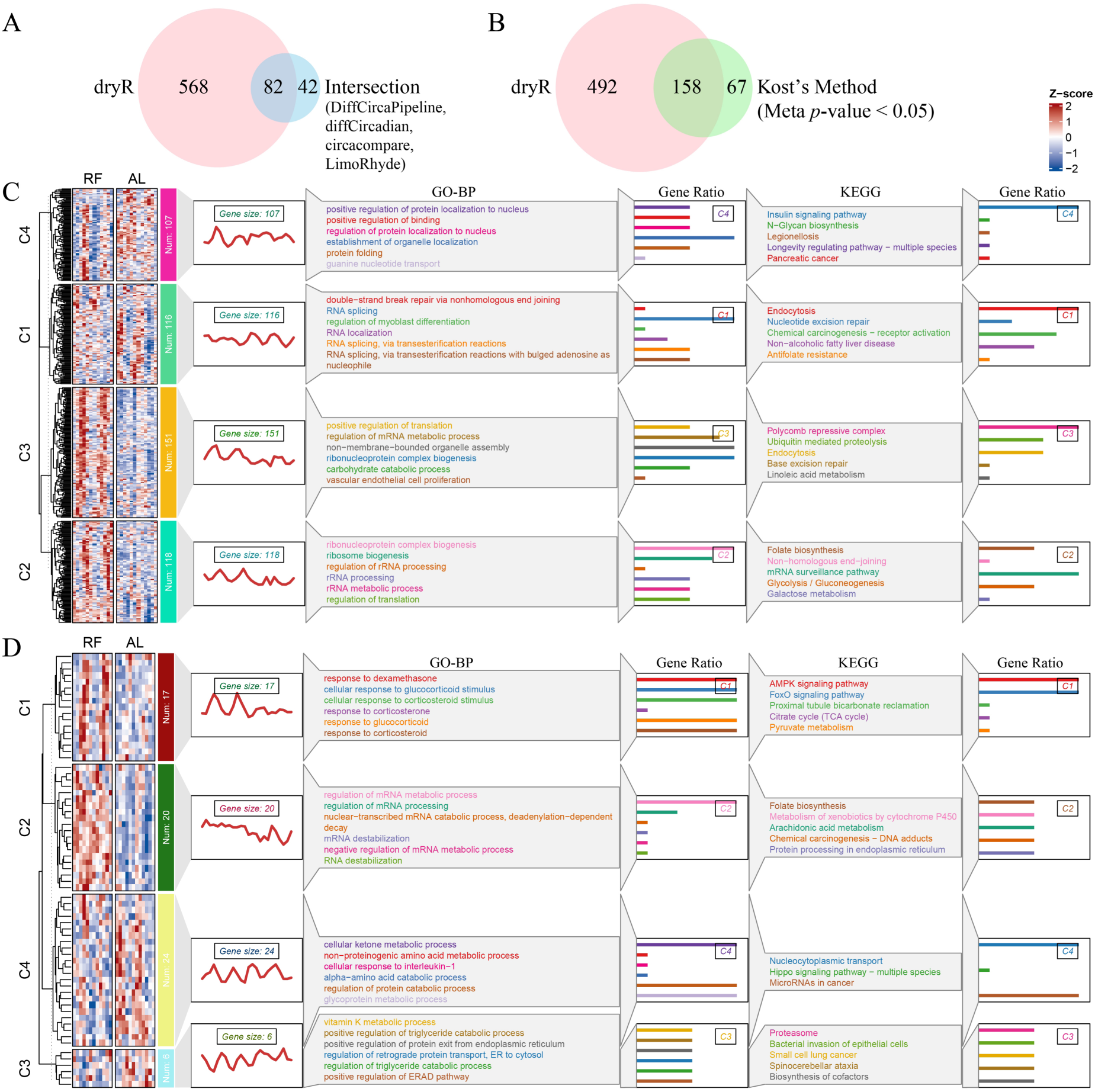
Validation of the consensus framework on the liver feeding-rhythm dataset. Performance comparison among intersection consensus, DiffRhyShiny’s Meta *p*-value (Kost’s Method), and dryR using *Bmal1* KO ad libitum (AL) vs. night-restricted feeding (RF) transcriptomics. **(A– B)** Venn diagrams showing DRG recovery rates against dryR baseline for **(A)** intersection consensus (12.62%) and **(B)** Kost’s Method (24.31%). **(C–D)** Expression heatmaps (Z-score), trend lines, and top-ranked GO-BP/KEGG pathways for disjoint gene sets uniquely identified by **(C)** dryR or **(D)** Kost’s Method. Alt text: Venn diagrams, gene expression heatmaps, and trend lines comparing the recovery rates and functional enrichment of differentially rhythmic genes identified by intersection consensus, dryR, and Kost’s Method in an independent mouse liver dataset.

## Discussion

The rapid development of differential rhythmicity algorithms has created a need for systematic evaluation of their performance under diverse analytical conditions. In this study, we established a benchmarking framework that evaluated seven representative differential rhythmicity methods across four dimensions: classification accuracy under predefined rhythmic scenarios, detection of specific rhythmic parameter changes, robustness to noise differences between conditions, and the influence of sampling design factors including sampling density, observation duration, biological replication, and missing observations. A previous systematic comparison using the same *Bmal1* knockout dataset examined differences in threshold sensitivity and gene-level overlap among algorithms and demonstrated that methods generated distinct candidate gene sets because they define differential rhythmicity using different statistical criteria (Miao *et al*. 2024). Our study extends these observations by evaluating algorithm performance using predefined ground-truth scenarios. For example, when both conditions were rhythmic with identical parameters, compareRhythms (Pelikan *et al*. 2022) and dryR (Weger *et al*. 2021) achieved 86.1% classification accuracy, whereas diffCircadian (Ding *et al*. 2021) achieved 54.7% accuracy, representing a 31.4 percentage-point difference under this simulation condition (Fig. 3C). We also observed non-monotonic power patterns for DODR (Thaben and Westermark 2016) and LimoRhyde (Singer and Hughey 2019, Obodo, Outland, and Hughey 2023) under specific noise heterogeneity scenarios (Fig. 5B), indicating that algorithm performance can vary when groups differ in signal-to-noise characteristics. Furthermore, our sampling simulations showed that total observation number had a stronger influence on classification accuracy than the distribution between sampling frequency and observation duration under the evaluated simulation conditions (Fig. 6).

Across the evaluated scenarios, algorithm performance was strongly influenced by whether the statistical model incorporated the rhythmic features being tested. Methods that do not explicitly evaluate specific rhythmic parameters may have limited ability to detect corresponding changes under certain simulation conditions. For example, algorithms without mesor-related testing procedures showed reduced power for detecting mesor alterations, whereas methods incorporating mesor estimation achieved higher detection performance in this scenario (Fig. 4D). Similarly, algorithms incorporating model-fit information showed distinct responses under noise heterogeneity simulations compared with methods primarily focused on rhythmic parameter differences (Fig. 5B). These findings suggest that algorithm selection should be guided by the expected biological alteration and the statistical features modeled by each method. Although some algorithms showed advantages in specific scenarios, no single method achieved the highest performance across all evaluated conditions. Therefore, consensus approaches may provide complementary information when the underlying biological mechanism involves multiple aspects of rhythmic regulation.

The second major contribution of this study is the integration of benchmarking results into an accessible computational platform. DiffRhyShiny combines seven differential rhythmicity algorithms through a unified data interface, incorporates Kost’s Method for dependent *p*-value integration (Kost and McDermott 2002), and provides interactive visualization of individual and consensus results (Fig. 7). To our knowledge, this represents an application of dependence-aware *p*-value combination to differential rhythmicity analysis. This approach is particularly relevant because multiple algorithms analyze the same expression profiles and therefore generate correlated statistical evidence. Unlike Fisher’s method, which assumes independence among combined tests, Kost’s Method incorporates covariance information among *p*-values during statistical integration. In our simulation framework, Kost’s Method maintained Type I error rates close to the nominal level under correlated evidence structures (Fig. 8A) and showed improved detection performance compared with more conservative combination strategies under several alternative scenarios (Fig. 8B-D). These results suggest that dependence-aware evidence integration can provide an alternative to strict intersection-based approaches, which may exclude genes supported by complementary statistical evidence.

DiffRhyShiny complements existing circadian analysis resources by focusing specifically on differential rhythmicity benchmarking and multi-method integration. Previous tools, including DiffCircaPipeline’s Shiny application and CosinorTest, provide interactive analysis environments for specific circadian modeling approaches (Xue *et al*. 2023, Ding *et al*. 2024). Other resources have focused primarily on single-condition rhythmicity detection rather than systematic differential rhythmicity comparison (Wu B and Ja 2025, Han *et al*. 2026). DiffRhyShiny extends these efforts by combining algorithm benchmarking, dependence-aware consensus analysis, and interactive visualization within a unified workflow. By providing access to multiple analytical strategies within the same platform, DiffRhyShiny allows researchers to compare individual algorithm outputs and evaluate integrated evidence without requiring extensive programming experience.

Several limitations should be considered when interpreting this study. First, although DiffRhyShiny was evaluated using two biological contexts, including *Bmal1* knockout fibroblast transcriptomes and an independent liver feeding-rhythm dataset, additional datasets involving different species, tissues, and experimental perturbations will be required to further assess generalizability. Second, the cloud-based deployment of DiffRhyShiny may encounter computational limitations for extremely large datasets because of shared computing resources; local deployment may therefore be preferable for large-scale analyses. Third, our benchmarking primarily used cosine-based generative models, which represent assumptions shared by several circadian analysis methods but do not capture all possible waveform structures, including asymmetric rhythms and higher harmonics. Future benchmarking studies should incorporate diverse waveform models and emerging datasets, including single-cell time-series experiments where sampling constraints and biological variability present additional challenges (Xu *et al*. 2024).

### Conclusion

Differential rhythmicity analysis is increasingly used to investigate condition-specific changes in biological oscillations, but the diversity of available algorithms creates challenges in method selection and result interpretation. In this study, we systematically benchmarked seven differential rhythmicity algorithms across multiple simulation scenarios and transcriptomic datasets, demonstrating that algorithm performance depends on the rhythmic features being evaluated and the characteristics of the experimental design. To integrate complementary evidence from multiple methods, we introduced a Kost’s Method-based consensus framework that accounts for dependency among statistical tests and implemented this approach in DiffRhyShiny, an interactive platform for differential rhythmicity analysis.

DiffRhyShiny provides a unified environment for algorithm comparison, consensus analysis, and visualization, enabling researchers to examine differential rhythmicity from multiple statistical perspectives within a single workflow. By combining systematic benchmarking with dependence-aware evidence integration, DiffRhyShiny provides a standardized framework for selecting, comparing, and integrating differential rhythmicity methods across diverse circadian transcriptomic studies.

## Data availability

The publicly available transcriptomic datasets analyzed in this study are deposited in the Gene Expression Omnibus (GEO) repository. Specifically, the mouse fibroblast RNA-seq dataset is available under accession number GSE253826 (https://www.ncbi.nlm.nih.gov/geo/query/acc.cgi?acc=GSE253826) (Unruh *et al*. 2024), and the independent liver feeding-rhythm dataset used for biological validation is available under accession number GSE135898 (https://www.ncbi.nlm.nih.gov/geo/query/acc.cgi?acc=GSE135898) (Weger *et al*. 2021). The DiffRhyShiny interactive web platform developed in this study is freely accessible at https://omicsai.shinyapps.io/diffrhyshiny. The open-source code of the web platform, which allows users to deploy the tool on their local computers or servers, is available on GitHub at https://github.com/liuchuan111/diffrhyshiny. The code and data to reproduce all benchmarking results are archived on Figshare at https://doi.org/10.6084/m9.figshare.33070100 (Liu *et al*. 2026).

## Author Contributions

**Chuan Liu:** Conceptualization, Methodology, Software, Writing -Original Draft, Writing -Review & Editing, Funding acquisition. **Lu Liu:** Software, Formal analysis, Investigation, Writing -Original Draft. **Na Li:** Investigation, Project administration, Funding acquisition. **Yang Jiang:** Data Curation, Formal analysis. **Chunyan Li:** Data Curation, Formal analysis. **Ying Liu:** Formal analysis. **Yongan Fu:** Data Curation. **Kai Guo:** Conceptualization, Writing -Review & Editing, Supervision, Funding acquisition. All authors read and approved the final manuscript.

## Funding

This work was supported by the Scientific and Technological Research Program of Chongqing Municipal Education Commission [grant numbers KJQN202600615, KJQN202300616] and the Chongqing Natural Science Foundation [grant number CSTB2023NSCQ-MSX0582].

## Conflict of interests

The authors declare no competing interests.

## Notes

### Competing Interest Statement

The authors have declared no competing interest.

